# *In situ* control of root–bacteria interactions using optical trapping in transparent soil

**DOI:** 10.1101/2021.12.18.473189

**Authors:** Sisi Ge, Kathryn M Wright, Sonia N Humphris, Lionel X Dupuy, Michael P MacDonald

## Abstract

Bacterial attachment on root surfaces is an important step preceding the colonisation or internalisation and subsequent infection of plants by pathogens. Unfortunately, bacterial attachment is not well understood because the phenomenon is difficult to observe. Here we assessed whether this limitation could be overcome using optical trapping approaches. We have developed a system based on counter-propagating beams and studied its ability to guide *Pectobacterium atrosepticum* (Pba) cells to different root cell types within the interstices of transparent soils. Bacterial cells were successfully trapped and guided to root hair cells, epidermis cells, border cells and tissues damaged by laser ablation. Finally, we used the system to quantify the bacterial cell detachment rate of Pba cells on root surfaces following reversible attachment. Optical trapping techniques could greatly enhance our ability to deterministically characterise mechanisms linked to attachment and formation of biofilms in the rhizosphere.

## Introduction

Crop and soil bacteria depend heavily on each other to acquire nutrients and grow healthily. Bacteria benefit from nutrients deposited in soil by plant roots in the form of exudates, mucilage, and border cells [1]. They sense a huge diversity of root-derived compounds [2], move chemotactically towards these sources of nutrient [3] and colonise the soil volume where these resources are abundant [4]. Many species of bacteria, in turn, contribute to essential soil functions, enabling bioavailability of nutrients to plant roots by, e.g. breaking down organic compounds [5], solubilising mineral ions [6] or fixing nitrogen [7]. Other bacteria can be pathogenic and contribute to the spread of diseases with a significant economic impact on farming [8, 9].

The broad range of interactions occurring in soil between plants and microbes creates a complex ecosystem termed the rhizosphere. Microbial biodiversity in the rhizosphere has been well documented [10, 11], but its dynamics remains mysterious because mechanistic knowledge of the interactions between microbes and plant cells are poorly understood. Rhizobacteria are known to successfully attach to the root surface and subsequently form biofilms [12, 13], but the process depends largely on the plant species colonised [14] and is facilitated by multiple bacterial processes the genes for which are now being uncovered [15]. The ability of bacteria to colonise root surfaces and to access root-derived nutrients, or cause disease, also depends on a number of plant ecophysiological factors [16]. For instance, the ability of phytopathogens to cause infection is believed to be influenced by the presence of lesions on the root [17, 18] and the colonisation of plant growth promoting rhizobacteria is known to be affected by competition with other microorganisms [19].

Successful colonisation of root surfaces is initiated by contact between a bacterium and a plant cell [20]. Unfortunately, the tools available to understand the conditions for how such contacts lead to attachment and colonisation are limited. Colonisation assays are commonly used to determine proliferation on root cells [21, 22] and molecular approaches can also be used to determine microbial diversity [23]. Quantification of plant-bacteria contacts is not, to date, attainable, and determination of the attachment and proliferation rate is indirect [24]. Live microscopy can be used to track the activity of a single bacterium and observe contacts with plant cell walls [25]. However, microscopes with sufficient resolution to track an individual bacterium typically lack field-of-view, or the inherent stochasticity of bacterial movement makes data collection inefficient.

The development of Transparent Soils (TS) [26, 27] has opened new avenues for direct manipulation of individual bacterial cells *in situ* using optical trapping. Transparent soils create a porous structure and microhabitats for the establishment of rhizosphere properties. Because the soil particles are transparent, live observation and optical manipulation are enabled. A combination of transparent soil with optical trapping of rhizobacteria *in situ* could greatly enhance our ability to control the number and location of bacteria attaching to plant roots and facilitate long term observations of the maintenance, and subsequent proliferation, of bacteria on root surfaces. Because optical trapping enables manipulation of single cells, bacteria-root attachment can be done in a controlled and deterministic way, allowing for an increase in experimental throughput, where we know both where *and* when the root-bacterial interactions will take place.

The objective of this study is to develop a system for optical trapping and guiding of single or multiple bacterial cells through transparent soil. The system developed is based on counter-propagating laser beams, which we combined with an imaging and pulsed laser dissection system [28] to track the movement of bacteria following attachment on healthy tissue and on lesions. We optimised the optical parameters to manipulate bacterial cells on root surfaces and studied the factors affecting the attachment of the pathogenic bacteria *Pectobacterium atrosepticum* (Pba).

## Material and methods

### Optical trapping instrument

A dual-beam counterpropagating trapping set up, similar to Ashkin’s original arrangement [29], was chosen to allow for optical manipulation with large working distances and access from two sides of the sample chamber to allow guiding from either side. A CW 1064 nm Ytterbium fibre laser (PYL-2-1064-LP, IPG Laser GmbH, Germany) with maximum output power of 2 W, and a beam diameter at the output of 5 mm, was used to build the optical trapping instrument (Figure 1). The beam was aligned and expanded using a telescope made of plano-convex lenses (LA4148-C, LA4380-C, Thorlabs, UK). A 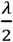 waveplate (WPH05M-1064, Thorlabs, UK) changed the portions of vertical and horizontal polarisations of the laser beam, and a polarising beam splitter (CCM1-PBS253/M, Thorlabs, UK) then split the beam into two illumination arms with different polarisations. The combinations of the 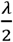 waveplate and polarised beam splitter were used to adjust the laser power of each illumination arm independently, and to ensure the counterpropagating beams do not interfere through the imposition of orthogonal polarisations. The laser beam was then directed through a series of mirrors and a 4-f image relay system (L_1_ / L_2_ and L_3_ / L_4_ respectively LA4380-C and LA4874-C, Thorlabs, UK), which allowed the adjustment of the laser beam vertically or horizontally without tilt and or loss of collimation. Two aspherical lenses (C240TMD-C, f = 8.00 mm, NA=0.5, Thorlabs, UK) were used for focusing the laser beam into the soil sample. Samples chambers were anchored on a 3-axis linear translation stage (M-562-XYZ, Newport, UK) with 13 mm of travel in each direction.

**Figure 1.**
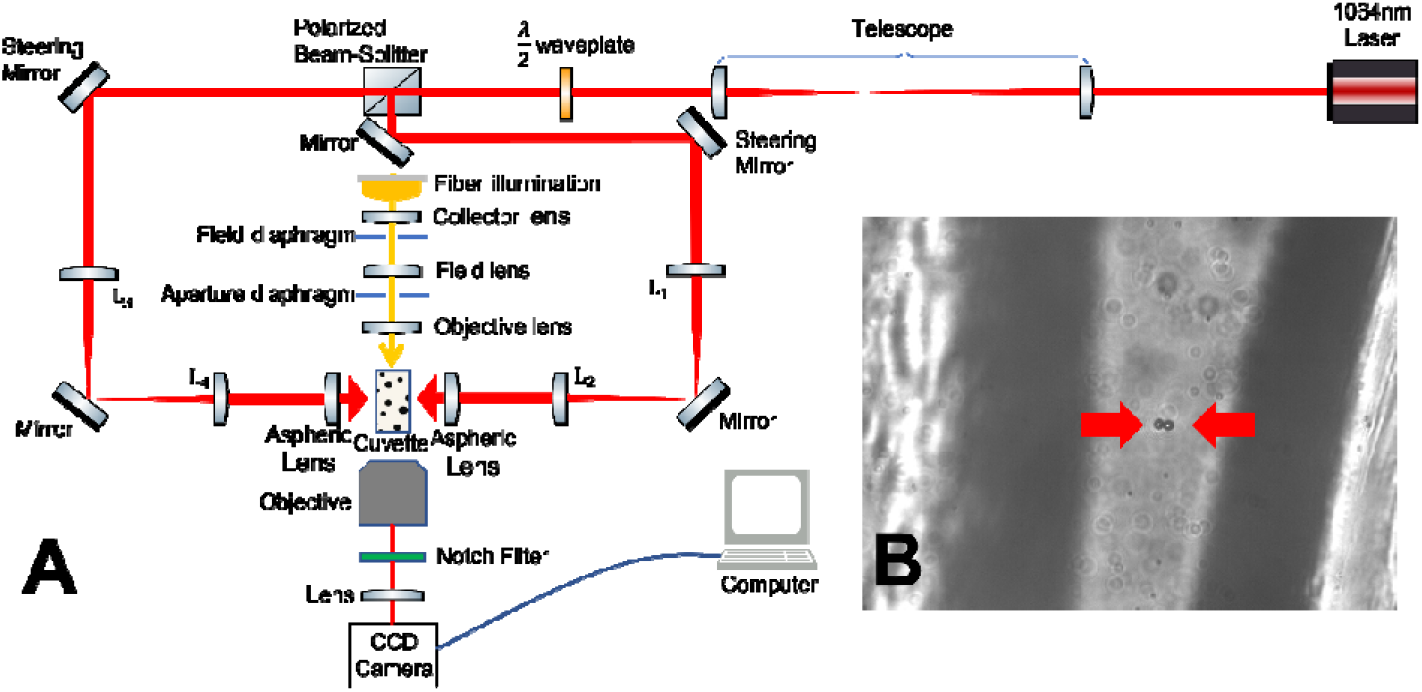
Set-up for trapping and guiding bacterial cells in transparent soil. (A) A system with counter-propagating laser beams was developed to perform optical trapping in transparent soil. A 1064 nm near-infrared laser was expanded using a telescope. A /2 waveplate and a polarising beam-splitter were used to control the laser power for each arm. The laser beams subsequently went through a 4-f system in which the steering mirror and the back aperture of an aspheric lens are conjugated. The construction ensures any laser angular movement from the steering mirror translates into lateral movement of the beam, preventing the deformation of the focal plane. The imaging arm included a notch filter for filtering of scattered light and a CCD camera acquiring image data projected from a Köhler illumination system. (B) Two polymer beads (red arrows) trapped between transparent soil particles.

The imaging arm consisted of an objective (M Plan Apo SL 20X, M Plan Apo 50X, M Plan Apo 100X, Mitutoyo, UK), a notch filter that attenuated the scattered trapping light (NF1064-44, CWL = 1064 ± 2 nm, FWHM = 44 nm, Thorlabs, UK), a tube lens (TTL200-A, Thorlabs, UK), and a progressive scan CMOS camera sensor (Grasshopper3 GS3-U3-23S6M, FLIR Systems, USA). Bright-field images were obtained using a Köhler illumination system assembled from a fibre illuminator (HPLS200, Thorlabs, UK), lenses (LA1422-A, LA1131-A, LA1509-A, Thorlabs, UK) and iris diaphragms (SM2D25D, Thorlabs, UK). The power used to hold particles was measured directly using a power meter (PM100D, Thorlabs, UK) and a photodiode power sensor (S121C, Thorlabs, UK). Distance measurements within the sample were calibrated using a reticle (horizontal micrometre scales transmission reticles, Edmund Optics, UK).

### Laser ablation

Laser ablation was performed using a different system [28]. A 532 nm Q-switched pulsed laser source (Continuum minilite II) and an aspherical lens (C280TMD-C, f = 18.40 mm, NA = 0.15) were used to focus the beam on the sample. The laser provided high peak power of 5.0 × 10^6^ W to 8.3 × 10^6^ W in pulses of nanoseconds in duration, delivering precise dissection with minimal thermal damage to the surrounding root tissues. The aspherical lens of the ablation instrument was chosen to achieve a small focal spot at the required working distance in a lens, whilst also remaining tolerant of the high peak powers of the laser pulses. Targeting of the ablation was achieved by moving root samples anchored on the translation stage. The imaging system consisted of a Köhler illumination, an imaging objective (TU Plan ELWD 20X, Nikon, UK), a notch filter (NF533-17, Thorlabs, UK), a tube lens (LA1509-A, Thorlabs, UK) and a CCD camera (Guppy F-046, Allied Vision, Germany) acquiring images at 10 frames per second.

### Sample preparation

Transparent soil was prepared from Nafion® pellets (Ion Power Inc., USA). The pellets were fractured using a cryogenic mill (SPEX SamplePrep 6770) and milled to obtain particle sizes of between 250 µm and 1,250 µm in size. The resulting Nafion particles were converted to anionic form by a series of chemical treatments described in Downie [26]. Fractured Nafion® particles were titrated using 4.4g L^-1^ Murashige and Skoog Basal Medium (Sigma M5519) and autoclaved for 20 minutes at 121 °C. Soil samples were prepared in standard spectrophotometer cuvettes (3/G/5, Starna Scientific, UK). Water was added to the sample to approximately match the refractive index of the Nafion particles. Additionally, the cuvettes were UV sterilised (ELC-500, 365nm, Fusionet LLC, USA) to minimise the risk of biological contamination before inoculation with bacterial suspension or seedlings (Visualization 1). Cuvettes were sealed with parafilm tape (PM996, Cole-Parmer, UK) before use in optical trapping and guiding experiments.

*Pectobacterium atrosepticum* (Pba) is an important plant pathogen that is known to infect potato plants and cause tuber rot [30]. It is a rod-shaped bacterium with a width of between 0.5 µm and 1.0 µm and a length of between 1.0 µm and 3.0 µm, a size range known to work well in optical trapping. Preliminary tests of optical manipulation instruments were performed using polymer beads that match the size, if not the shape and refractive index, of the bacterium (1 µm, 2 µm polymer beads 4010A, 4202A, ThermoFisher Scientific, UK). The particles were diluted to 2.1 × 10^−3^ g ml^-1^ before transfer to glass cuvette with or without transparent soil. Cell suspensions of *Pectobacterium atrosepticum* SCRI1039 (Pba) were cultured overnight with aeration in Lysogeny broth at 27°C, harvested by centrifugation (4000 rpm, 10 minutes, 18°C) and the pellet resuspended in 10 mM MgSO_4_. The cells were re-sedimented and washed again before being resuspended in 10 mM MgSO_4_ at a final concentration of 10^7^ CFU ml^-1^. Prior to imaging and optical trapping experiments, Pba suspensions were diluted in distilled water to 8 ×10^4^ CFU ml^-1^.

Lettuce seeds (*Lactuca sativa*) were surface sterilised in 10% bleach (Domestos, UK) for 20 minutes following multiple washes in distilled water. Seeds were germinated in microbiology grade agar prepared at 1.5% w/v in distilled water overnight and subsequently transferred to glass cuvettes containing transparent soil. Plants were then grown in cuvettes containing water and transparent soil at room temperature (approximately 20 °C) for 24 to 48 hours before inoculation.

### Experiment 1 – characterisation of the trapping

Preliminary experiments were performed to determine the minimum power needed to trap and hold a particle within the laser beam. The minimum trapping power represents the smallest power output needed to directly trap a particle in the focus of the beam. The minimum holding power was measured by trapping a particle at a higher power first and then decreasing the power output of the laser beam, which was defined as the smallest power output needed to keep the particle in trap. Polymer beads and bacteria trapping in both water and transparent soil were recorded as supplement (Visualization 2 to 5).

To determine the speed at which particles are attracted towards the counter-propagating beams, both polymer beads and Pba suspensions were prepared in glass cuvettes. A separate sample cuvette was prepared for each type of particle (1 µm polymer beads, 2 µm polymer beads and cell suspensions of Pba) and for each medium (water or transparent soil). The cuvettes were in turn fixed on the stage of the optical trapping instrument, and the position of the sample was adjusted using a 3-axis linear translation stage. The laser beam was subsequently applied for 45 s, the movement of particles induced by the beam was recorded by the imaging system, and the velocity of particles was measured from image data. The measurement was recorded three times at a given position, with an interval of 30 s between measurements, before moving to a new location. Three different locations were studied at any given power output of the laser beam. When experiments were performed in transparent soil, measurements were taken at four different locations and repeated four times at each location. The laser was operated at a power of 150 mW, 200 mW, 250 mW and 300 mW (measured at the focal plane of the aspherical lens).

In a third step, guiding numbers and speed were studied on live samples. Lettuce seedlings were grown in transparent soil, and 200 μL of Pba cells suspension (8 × 10^4^ cells per millilitre) were introduced into the cuvette using a single channel pipette before experiments. Samples were left on the translation stage for 5 to 20 minutes for the bacteria to disperse in the soil solution and be observable in the root area. In order to control the number of bacterial cells guided to the surface of the root, we studied how the duration of the application of the laser light affected the number of cells attached to the root surface. The laser beam was first positioned at a location on the surface of the root epidermis. In this case, only one beam was used with the increased power output of 400 mW for durations of 1 s, 4 s, 10 s, 20 s and 30 s. Live image data was acquired to count the number of bacterial cells attracted to the root surface. For each duration, data was recorded at three different locations each, from three different samples.

### Experiment 2 – guiding bacteria to different root cell types

Pba bacterial cells were guided to root cell types that could easily be identified by live imaging. These include epidermis cells, border cells, root hair cells and cells that have been damaged by laser dissection. To guide bacterial cells towards root lesions, roots were first ablated using the laser dissection instrument before transfer to the optical trapping instrument for the remainder of the experiment.

In these experiments, all cuvettes containing samples were placed on the sample stage of the optical trapping instrument, and live images captured by the camera were used to locate root cells of interest. Even though only one arm of the laser was used for guiding bacterial cells; having two beams allow guiding from either side of the sample depending on where a blockage might be. The power output of the trapping laser beam was set to 400 mW at the focal plane of the aspherical lens with less than 50% of the power reaching the focal plane. Bacteria were guided to the surface of the root by the laser beam, and the movement of groups of bacteria on the surface of the root was caused by the translation of the stage. In total, 40 locations were monitored from 5 samples.

### Experiment 3 – quantification of cell detachment rate from epidermis cells

In the final series of experiments, we used the optical trapping system to facilitate quantification of bacterial cell attachment and identify factors affecting the cell detachment rate. Between five and fifteen bacterial cells were first guided to the surface of the root epidermis using laser application times of between 5 s and 10 s. The cells were spread along the root surface to ensure all bacteria were in contact with the epidermis. The laser beam was subsequently switched off, Pba cells were allowed to detach and disperse in the medium and the number of cells detaching from the surface of the root was recorded for 30 seconds after the laser beam was switched off. We subsequently analysed the live image data to quantify how the cell detachment rate changed as a function of time and the number of cells. In total, we followed the detachment of Pba cells from the root epidermis at 10 different locations from three samples. Similar experiments were performed using 1 µm polymer beads guided to and spread on root epidermis. However, as polymer beads tend to detach more rapidly, it was difficult to count the number of beads with certainty. In this case, the detachment rate measured is approximate and must be read as an estimate only of the lower bound of the true detachment rate.

Experiments were also performed to assess whether the presence of lesions in the plant tissue and bacterial cell agglomeration affect the cell detachment rate of bacteria. 8 lesions were produced by laser ablation of roots from 3 samples. Out of the 8 lesions, bacteria were guided to the root and spread on four of these lesions. Bacteria were guided agglomerated to the root without spreading on the remaining four lesions. A further 4 samples did not contain lesions. On four locations from two of these samples, bacteria were guided to the root epidermis and left agglomerated, and on four locations from the remaining two samples, bacterial cells were guided and spread on the epidermis.

Live brightfield imaging was used to determine the cell detachment rate. In each optical trapping experiment, a group (*N*_*tot*_) of bacteria was formed on the surface of the root. The number of detached bacteria (*N*_*det*_) was measured at a time *t* following interruption of the laser beam. The bacterial cell detachment rate was then calculated as

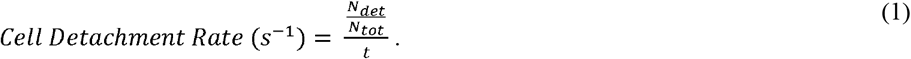

The cell detachment rate, therefore, indicates the fraction of the initial bacterial number detaching per second.

### Data analysis

Measurements of bacterial cell detachment rate were performed from live image data. Image data were analysed using Fiji [31]. In order to quantify the movement of cells and count the number of detached bacteria, we used the cell counter function [32]. All data is presented as mean ± standard deviation.

## Results

### Speed and number of particles trapped by the laser beam

The application of the trapping laser induced particles to move towards the focus of the beam, and the results showed that both single and multiple particles can be trapped by the beam (Figure 2A). The minimum power output of the laser beam required to trap a 1 µm polymer beads was (19 ± 6) mW. The minimum power output of the laser beam required to trap a 2 µm polymer beads was (63 ± 8) mW and for Pba it was (135 ± 7) mW, and three power readings were taken for each sample’s measurement.

**Figure 2.**
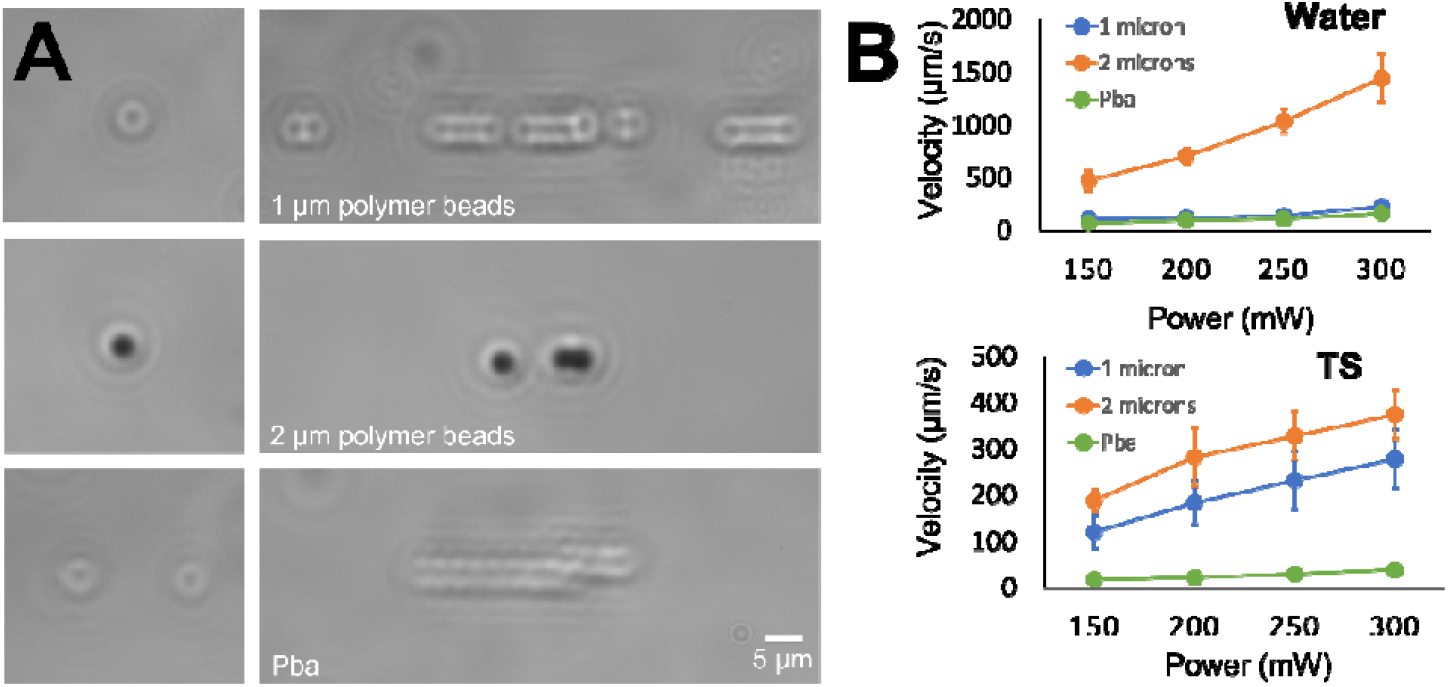
Guiding velocity in water and transparent soil (TS). A - Tests consisted of polymer beads and Pectobacterium atrosepticum cells in suspension in water or transparent soil. The velocity of both single particles (left) and multiple particles (right) was observed as a function of power output of the laser beam. The particles tested include 1 µm particles (top), 2 µm particles (middle) and Pectobacterium atrosepticum cells (bottom). B - When the power output of the laser beam increased the speed at which particles could be guided both in water and transparent soil also increased. The velocity observed in water (top) was consistently higher than the velocity recorded in transparent soil (bottom) for 2 µm particles and Pba cells, while the velocity of 1 µm particles in transparent soil was slightly higher than in water. Transparent soil caused the velocity of the guided 1 µm particles to be more variable but was lower for 2 µm particles and similar for the Pba cells.

The apparatus developed allowed vertical and horizontal adjustment of particles on the root surface by translating the sample. When the laser beam trapped the particles, controlled displacement was successfully achieved across the entire field of view of the imaging system (± 130 µm from the centre).

The average velocity achieved when guiding particles and Pba in water increased with the power output of the laser beam. Similarly, velocity of particles in transparent soil increases with the power output of the laser beam (Figure 2B, Table 1). The velocity for each sample at each laser power was measured from 5 different locations.

**Table 1:** Guiding velocity of polymer beads and Pba (μm/s)

| Table 1: Guiding velocity of polymer beads and Pba ($\mu\text{m/s}$) | | | | | | | | |
| --- | --- | --- | --- | --- | --- | --- | --- | --- |
| Sample \ Power | Water |  |  |  | Transparent Soil |  |  |  |
|  | 150 mW | 200 mW | 250 mW | 300 mW | 150 mW | 200 mW | 250 mW | 300 mW |
| 1 $\mu\text{m}$ beads | 110 ± 20 | 120 ± 30 | 140 ± 20 | 230 ± 20 | 120 ± 40 | 180 ± 50 | 230 ± 60 | 280 ± 60 |
| 2 $\mu\text{m}$ beads | 480 ± 90 | 710 ± 70 | 1040 ± 120 | 1400 ± 200 | 190 ± 20 | 280 ± 60 | 330 ± 50 | 380 ± 50 |
| Pba | 73 ± 3 | 101 ± 17 | 113 ± 6 | 167 ± 7 | 19 ± 6 | 23 ± 6 | 31 ± 7 | 39 ± 9 |

### Guiding bacteria to root cells

The optical trapping system was used to guide bacterial cells and polymer beads to the root surface (epidermis cells) such that subsequent attachment rates could be measured, and results showed the number of cells attached to the root surface can be controlled via the duration over which the laser beam was applied (Figure 3). On average, bacteria were attracted to the surface of the root at a rate of (0.7 ± 0.1) s^-1^.

**Fig. 3.**
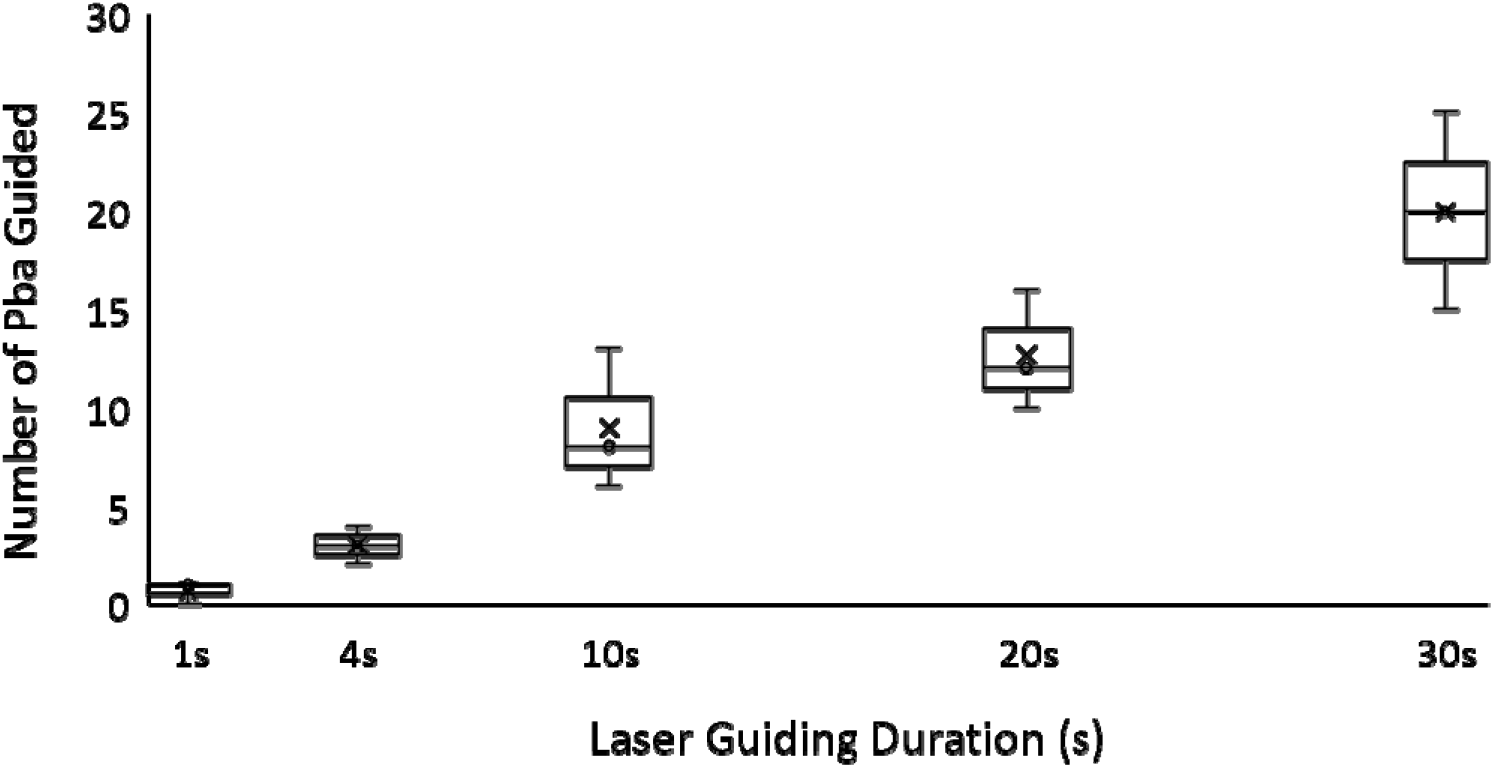
Optical trapping can be used for precise control of the number of bacteria attached to a specific location on the root surface. The number of Pba attached to the surface of epidermis cells was directly related to the time during which the laser was kept on. Therefore, the duration of the exposure to the laser beam can be used to control the number of cells attached to the root surface. The boxplot represents the variability of the data. Error bars of the boxplot indicate the 95% confidence interval, mean (cross) and median (diamond) value of the data.

The optical trapping system we developed could also be used to guide bacteria to other types of cells. Results showed that the optical trapping instrument could be used to move bacteria to root hair cells (Figure 4A), or border cells (Figure 4B). Although the laser beam often attracted multiple cells, results also showed that adjustment of the position of the beam and short exposure time could be used to manipulate single bacterial cells (Figure 4C). More complex experiments could also be performed using the system. These include the displacement of large groups of bacterial cells (Figure 5A) or inducing a lesion on the root tissue by laser ablation (Figure 5B) before guiding cells to the lesion (Figure 5C).

**Fig. 4.**
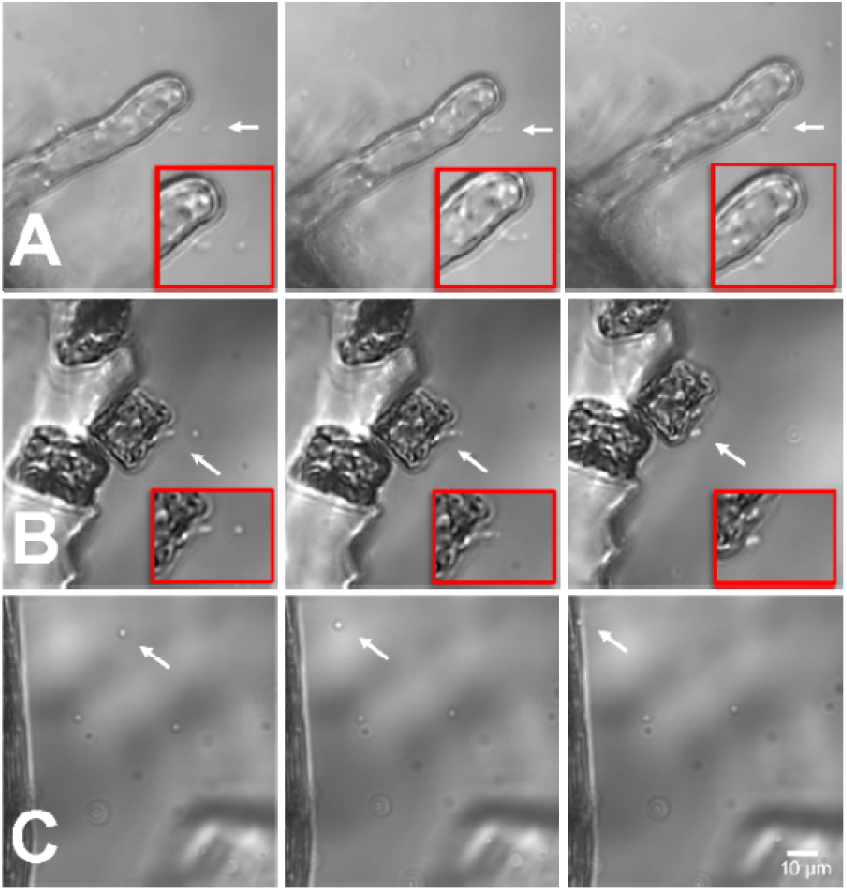
Time sequence showing the process of guiding of bacterial cells to specific root cell types. Guiding of bacteria was successfully performed on (A) root hair cells, B) border cells. C) and epidermis cells. Insets with red frames showed enlarged and clearer bacterial cells guiding events.

**Fig. 5.**
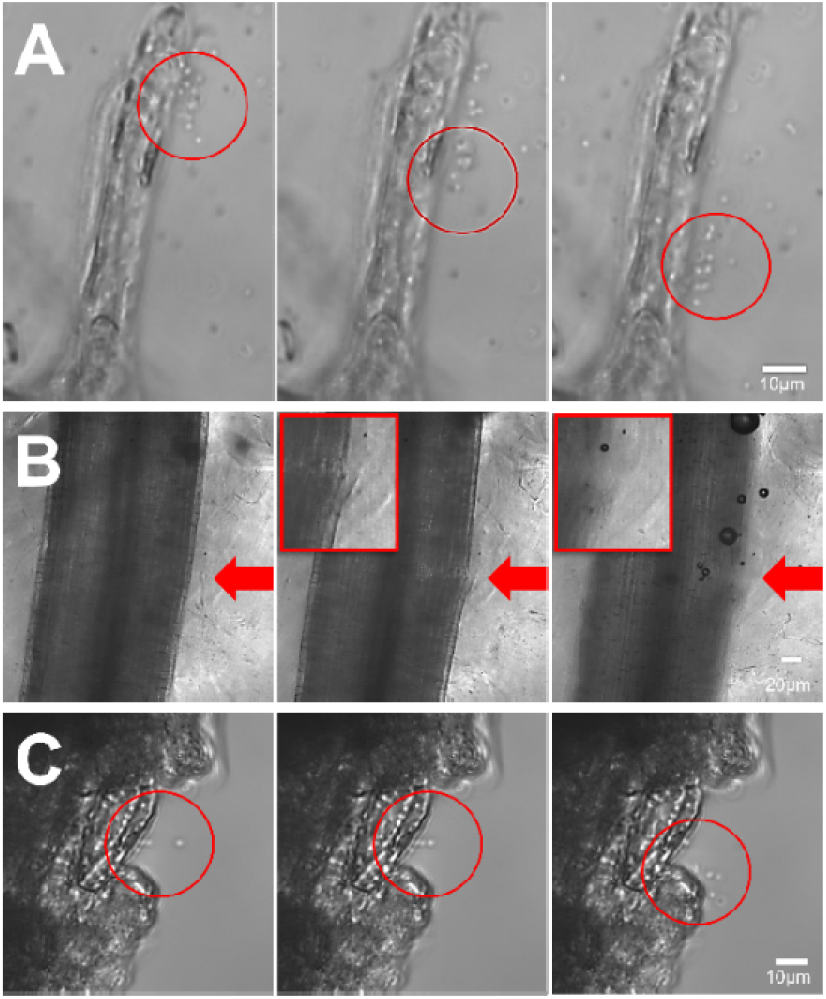
Optical trapping can also be used to perform manipulations such as (A) displacing an agglomeration of bacterial cells (circled in red), (B) performing laser ablation of the root tissue (insets with red frames showed enlarged and clearer images of root lesion and (C) guiding bacterial cell to the lesions induced by laser dissection (circled in red).

### Analysis of bacterial cell detachment from epidermis cells

We further investigated the factors that affect bacterial attachment on the root surface. The region of the root used for the experiments was free of bacterial cells at (t = 0), the laser was then switched on, and the number of bacterial cells observed on the surface of the root gradually increased (Figure 6A, left to right). After bacteria were attached, the laser was switched off, and the number of bacteria detaching from the root surface was recorded using time-lapse imaging (Figure 6B, SI video 6). Similar experiments performed using polymer beads produced significantly different results. The majority of the beads did not remain attached when the laser was turned off, making it difficult to quantify the detachment rate (SI video 7). Images were subsequently analysed to extract a dataset linking the number of bacteria on the surface of the root to the time after exposure to the guiding laser (Figure 6C).

**Fig. 6.**
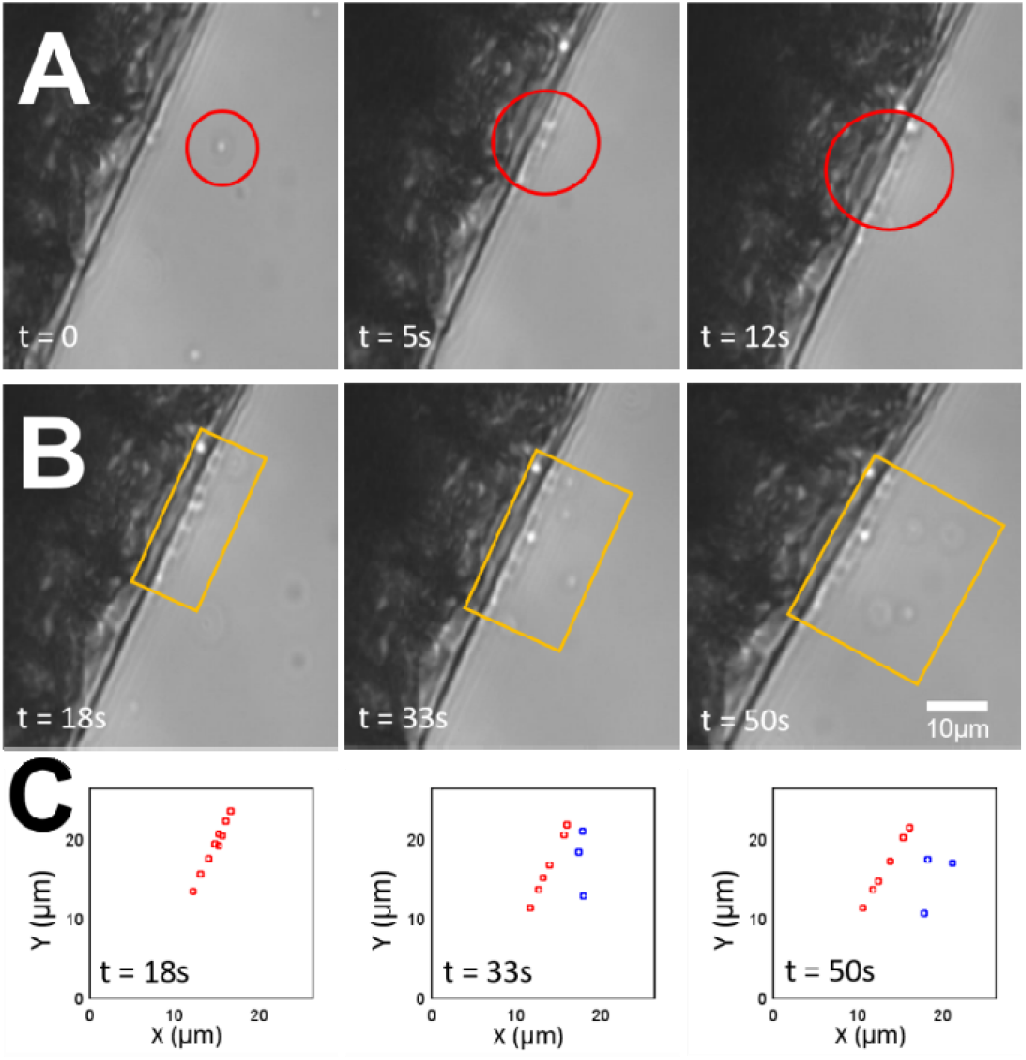
Using optical trapping to quantify bacterial cell detachment rate. (A) A group of bacterial cells is made to adhere to epidermis cells using optical trapping. The duration of application of the laser beam and the location of the focus is controlled so that bacterial cells are made to attach at the desired location and number (red circle). (B) The laser is subsequently switched off and the detachment of bacterial cells is monitored through time (yellow rectangle). (C) The videos of the detachment of bacterial cells are then used to track the cells and obtain quantitative information on the cell detachment rate of bacteria. The red symbols represent bacterial cells attached on the root surface and blue symbols represent detached bacterial cells, which derived from time-lapse images of (B).

The analysis of the data collected showed the average detachment rate from the epidermis cell was (0.07 ± 0.04) s^-1^. By comparison the detachment rate of polymer beads was two orders of magnitude larger [lower bound of (3 ± 1) s^-1^]. The detachment rate of Pba was also observed to decrease following the interruption of the laser. This trend was independent of the number of bacterial cells initially attached to the root (5 to 15 cells, Figure 7A). In the second step, we analysed whether the time a bacterium has been in contact with the root cell affects the cell detachment rate. Results from 36 cell detachment events showed there was a negative correlation between the bacterial cell detachment rate and the time a bacterium has remained attached (Figure 7B, R^2^ = 0.3796, p < 0.028).

**Fig. 7.**
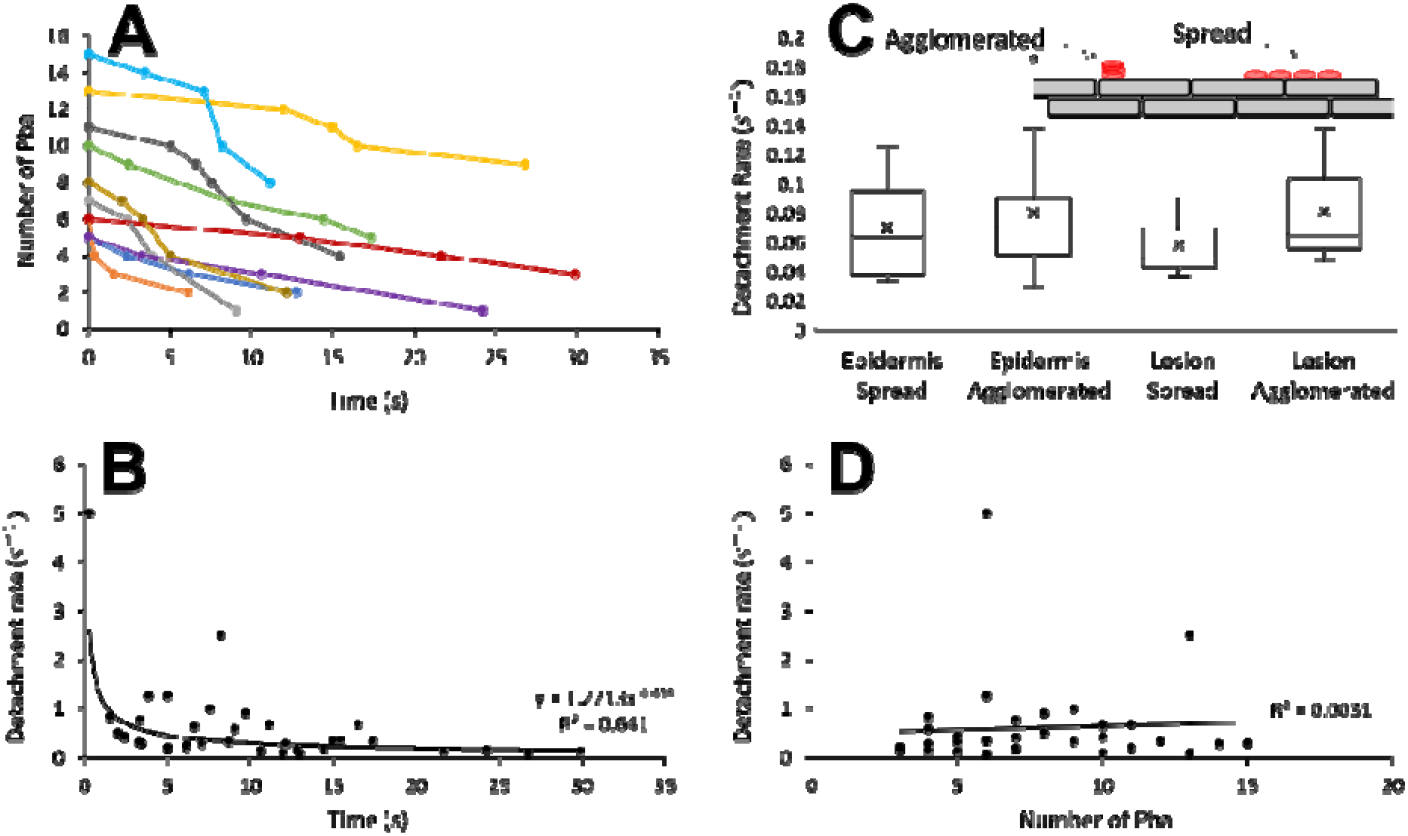
Factors affecting the persistence of attached bacterial cells on root epidermis. (A) A dataset was established to track how the number of bacterial cells decreases with time after the laser beam is switched off for 10 different observations. The dataset shows the number of bacteria remaining on the root epidermis declines consistently with time. (B) The decline of the number of bacterial cells on the epidermis slowed down with time, and correlation confirmed that the cell detachment rate is related to the time a cell adhered on the epidermis (exponential decline, p<0.028). (C) Further cell detachment experiments showed the number and location of bacterial cells do not significantly affect the cell detachment rate and agglomeration or lesions did not change the cell detachment rate (p = 0.346 and p = 0.866). (D) There was no observable correlation between the number of bacterial cells and the cell detachment rate (p = 0.746) when calculated from the decline in attached cell numbers shown in A.

We also studied whether the cell detachment rate was related to the agglomeration of cells, and the presence of a lesion on the root. Neither agglomeration nor lesion was observed to affect the cell detachment rate (based on the Kruskal – Wallis test, p = 0.284 and p = 0.937 respectively, Figure 7C). There was also no correlation observed between the cell detachment rate and the number of cells on the tissue calculated from the time-lapse data obtained in Figure 7A (p = 0.746, Figure 7D).

## Discussion

### Optical trapping of living cells in granular media

Optical trapping techniques have a decades long history of use for the three-dimensional manipulation of cells and organelles. The manipulation of single bacteria, viruses, cells and spores were demonstrated several decades ago [33-35]. Other applications also include the separation, sorting and rotation of cells [36-38] or unfolding of RNA structures [39]. Optical trapping has also been used to obtain quantitative measurements of forces and displacement of microscopic or nanoscopic structures. For example, the force and displacement of DNA strands were measured at piconewton and nanometre resolution [40, 41].

The application of optical trapping techniques to plant-microbe interactions is less common. Wang and co-workers [42] demonstrated the use of optical trapping for the levitation of single pollens and fungal spores. Optical trapping techniques have also been used to manipulate soil microorganisms that are not easily culturable such as nematodes [43] and bacteria [44]. The development of optical trapping in plant studies has largely focused on the manipulation of plant organelles. Using optical trapping, it is possible to manipulate organelles within the cytoplasm for the study of cytoplasm dynamics [45]. Optical trapping techniques were also used to immobilise subcellular structures within living cells and extend the time during which live microscopy can be performed [46]. The technique was also used in combination with Total Internal Reflection Fluorescence microscopy to study interactions between chloroplasts and peroxisomes [47]. More recently, optical trapping was successfully applied to induce controlled movement of amyloplasts in studies of root gravitropism [48].

### Requirements for optical manipulation of bacteria in the rhizosphere of transparent soils

The rhizosphere poses significant challenges to the application of optical trapping *in situ*. First, optical manipulation in natural soil is out of reach because of its opacity along with its structural and chemical complexity [49, 50]. Unlike in liquid or aeroponic cultures, soil substrates limit the movement of cells and particles and prevent the propagation of light [51, 52]. However, gels and aeroponics are not good models for the granular nature of real soils. The use of transparent soil facilitates optical manipulation inside soil-like, granular substrates. Because the roots grow through the complex soil structure of TS, optical trapping is made through an optically inhomogeneous medium. Optical aberrations occur when a laser passes through soil particles, and this can degrade the quality of the laser beam and may prevent the controlled displacement of cells [53]. Optical aberrations also exist in transparent soil as perfect refractive index matching between the soil particles and matching fluid, water in this instance, is never quite achieved. However, the transparent nature of the polymer soils used in this work mean that optical access into the rhizosphere is possible, with the question being whether the aberrations present degrade a trapping beam to the point of preventing optical manipulation deep within soil.

In this study, we have shown that moving bacterial cells within the inner structure of granular media using optical trapping is robust enough to work even without recourse to aberration correction techniques. The trapping and displacement of bacteria were achieved through water-filled interstices within the soil. The system we developed enabled precise displacement of a single or group of bacteria to the root surface (Figure 4-6). We demonstrated that the system can be used to quantify parameters for the attachment of bacteria on root cells (Figure 7).

During the experiments, optical manipulation was limited to a depth of 3 mm into the soil volume, up to which depth no aberration compensation was needed to achieve optical manipulation of polymer beads and bacteria. However, there is potential to improve the depth and precision of optical trapping techniques using more advanced optical aberration correction techniques. For example, digital micromirror arrays could be used to better control the size and shape of the optical trap [54], and the introduction of a spatial light modulator could allow the wavefront corrections needed to manipulate at depth, as was demonstrated for optical trapping through highly turbid media [55]. Optical manipulation could also be achieved using different beam types to adjust properties of the optical trap to the nature of samples or the medium used for growing the sample. Beam types include non-diffracting and “self-healing” Bessel beams [56], optical vortex beams [57] or Airy beams [58]. Furthermore, it has been shown that the combination of optimised beam types with aberration correction techniques can improve the properties of the trapping system [59, 60]. Acoustic [61] and magnetic tweezers [62] are also widely used micromanipulation techniques, but their application to soil and bacteria are limited. Acoustic trapping requires relatively uniform mass density and compressibility within the medium and this is difficult to achieve in a soil which is composed of gas, liquid and solid phases [63]. Magnetic micromanipulations are also problematic because bacterial cells are not superparamagnetic [64]. Both acoustic and magnetic tweezers have also limited spatial resolution and dexterity, making it difficult to manipulate a single bacterium.

### Opportunities for the study of root microbe interactions

Bacterial colonisation of the rhizosphere is largely dependent on bacteria attaching on surfaces and in turn forming biofilms [65]. Microbial attachment on root surfaces, however, is a complex process to observe. Bacterial cells are small particles which are subject to thermal effects [66, 67]. Their distribution in the soil volume is sparse [68]. Collision with a host does not consistently result in a cell binding with the host [69]. Both attractive and repulsive forces arise during a collision with a plant cell [20], and the plant immune response may also affect the ability of a microbe to establish on the root [70]. These experimental constraints make quantitative characterisation of the bacterial attachment process particularly elusive.

The work presented here showed that optical trapping can overcome these limitations, as the number of cells, their movement and site of attachment are controlled, experiments are not limited by the stochasticity of collisions with root surfaces and the observation of rare events. In our set-up the attachment of multiple cells could be performed on demand (within less than a minute), allowing deterministic characterisation studies of the factors affecting the bacterial detachment rate.

Although this study characterised only the properties of bacteria attachment on the surface of plant roots, there are now many opportunities to further study soil microbial processes in soil. Our instrument could be used, for example, to quantify attachment *in situ* and on various cell types, surfaces and at different stages of the formation of biofilms in the rhizosphere [71]. The energetics of bacterial mobility are largely unknown, yet remain an important determinant of microbial colonisation. Optical trapping could also be used to study the viscous forces experienced by bacteria when swimming in the pore space or quantify the surrounding viscosity induced by mucilage and exudation [72]. Attachment to other types of hosts or surfaces is also essential to the survival of many microbial species. Hence, with the help of techniques demonstrated, the great diversity of bacterial attachment mechanisms in soil could be explored, notably the attachment to soil particles [73], insects [74], earthworms [75].

## Funding

This work received funding from the Biotechnology and Biological Sciences Research Council (BB/T010657/1) and the European Research Council (ERC no 647857).

## Acknowledgement

We thank Daire Carroll for transparent soil preparation and plant samples, Gillian Fraser, Jacqueline Marshall and Lauren Watts for assistance with bacterial cultures.

## Disclosures

The authors declare no conflicts of interest.

## Data availability

Data underlying the results presented in this paper are not publicly available at this time but may be obtained from the authors upon reasonable request.

## Supplemental document

See visualization for supporting videos.

## Notes

### Competing Interest Statement

The authors have declared no competing interest.

## References

1. Fel Zahar Haichar, F., et al., Plant host habitat and root exudates shape soil bacterial community structure. The ISME journal, 2008. 2(12): p. 1221–1230.

2. Zhalnina, K., et al., Dynamic root exudate chemistry and microbial substrate preferences drive patterns in rhizosphere microbial community assembly. Nature Microbiology, 2018. 3(4): p. 470–480.

3. Allard-Massicotte, R., et al., Bacillus subtilis early colonization of Arabidopsis thaliana roots involves multiple chemotaxis receptors. MBio, 2016. 7(6): p. e01664–16.

4. Mercier, J. and S. Lindow, Role of leaf surface sugars in colonization of plants by bacterial epiphytes. Applied and Environmental Microbiology, 2000. 66(1): p. 369–374.

5. Gougoulias, C., J.M. Clark, and L.J. Shaw, The role of soil microbes in the global carbon cycle: tracking the below?ground microbial processing of plant?derived carbon for manipulating carbon dynamics in agricultural systems. Journal of the Science of Food and Agriculture, 2014. 94(12): p. 2362–2371.

6. Han, H.-S. and K. Lee, Effect of co-inoculation with phosphate and potassium solubilizing bacteria on mineral uptake and growth of pepper and cucumber. Plant Soil and Environment, 2006. 52(3): p. 130.

7. Zahran, H.H., Rhizobium-legume symbiosis and nitrogen fixation under severe conditions and in an arid climate. Microbiology and Molecular Biology Reviews, 1999. 63(4): p. 968–989.

8. Mansfield, J., et al., Top 10 plant pathogenic bacteria in molecular plant pathology. Molecular Plant Pathology, 2012. 13(6): p. 614–629.

9. Martins, P.M., et al., Persistence in phytopathogenic bacteria: do we know enough? Frontiers in Microbiology, 2018. 9: p. 1099.

10. Nihorimbere, V., et al., Beneficial effect of the rhizosphere microbial community for plant growth and health. Biotechnologie, Agronomie, Société et Environnement, 2011. 15(2): p. 327–337.

11. Kent, A.D. and E.W. Triplett, Microbial communities and their interactions in soil and rhizosphere ecosystems. Annual Reviews in Microbiology, 2002. 56(1): p. 211–236.

12. Bais, H.P., R. Fall, and J.M. Vivanco, Biocontrol of Bacillus subtilis against infection of Arabidopsis roots by Pseudomonas syringae is facilitated by biofilm formation and surfactin production. Plant Physiology, 2004. 134(1): p. 307–319.

13. Ramey, B.E., et al., Biofilm formation in plant–microbe associations. Current Opinion in Microbiology, 2004. 7(6): p. 602–609.

14. Fan, B., et al., Gram-positive rhizobacterium Bacillus amyloliquefaciens FZB42 colonizes three types of plants in different patterns. The Journal of Microbiology, 2012. 50(1): p. 38–44.

15. Knights, H.E., et al., Deciphering bacterial mechanisms of root colonization. Environmental Microbiology Reports, 2021.

16. Rodríguez-Navarro, D.N., M.S. Dardanelli, and J.E. Ruíz-Saínz, Attachment of bacteria to the roots of higher plants. FEMS Microbiology Letters, 2007. 272(2): p. 127–136.

17. Pfeilmeier, S., D.L. Caly, and J.G. Malone, Bacterial pathogenesis of plants: future challenges from a microbial perspective: challenges in bacterial molecular plant pathology. Molecular Plant Pathology, 2016. 17(8): p. 1298–1313.

18. Savatin, D.V., et al., Wounding in the plant tissue: the defense of a dangerous passage. Frontiers in Plant Science, 2014. 5: p. 470.

19. Benizri, E., E. Baudoin, and A. Guckert, Root colonization by inoculated plant growth-promoting rhizobacteria. Biocontrol Science and Technology, 2001. 11(5): p. 557–574.

20. Palmer, J., S. Flint, and J. Brooks, Bacterial cell attachment, the beginning of a biofilm. Journal of Industrial Microbiology and Biotechnology, 2007. 34(9): p. 577–588.

21. Gamalero, E., et al., Methods for studying root colonization by introduced beneficial bacteria. Sustainable Agriculture, 2009: p. 601–615.

22. Kloepper, J.W. and C.J. Beauchamp, A review of issues related to measuring colonization of plant roots by bacteria. Canadian journal of Microbiology, 1992. 38(12): p. 1219–1232.

23. Gollotte, A., D. Van Tuinen, and D. Atkinson, Diversity of arbuscular mycorrhizal fungi colonising roots of the grass species Agrostis capillaris and Lolium perenne in a field experiment. Mycorrhiza, 2004. 14(2): p. 111–117.

24. Carroll, D., et al., Framework for quantification of the dynamics of root colonization by Pseudomonas fluorescens isolate SBW25. Frontiers in Microbiology, 2020. 11: p. 2403.

25. Aldon, D., et al., A bacterial sensor of plant cell contact controls the transcriptional induction of Ralstonia solanacearum pathogenicity genes. The EMBO journal, 2000. 19(10): p. 2304–2314.

26. Downie, H., et al., Transparent soil for imaging the rhizosphere. PLoS One, 2012. 7(9): p. e44276.

27. Sharma, K., et al., Transparent soil microcosms for live-cell imaging and non-destructive stable isotope probing of soil microorganisms. Elife, 2020. 9: p. e56275.

28. Ge, S., L.X. Dupuy, and M.P. MacDonald, In situ laser manipulation of root tissues in transparent soil. Plant and Soil, 2021: p. 1–15.

29. Ashkin, A., Acceleration and trapping of particles by radiation pressure. Physical Review Letters, 1970. 24(4): p. 156.

30. Skelsey, P., et al., Threat of establishment of non-indigenous potato blackleg and tuber soft rot pathogens in Great Britain under climate change. PloS One, 2018. 13(10): p. e0205711.

31. Schindelin, J., et al., Fiji: an open-source platform for biological-image analysis. Nature Methods, 2012. 9(7): p. 676–682.

32. Rueden, C.T., De Vos, K. J., Guiet, R., Hiner, M. C., Cell_Counter. 2017.

33. Ashkin, A. and J.M. Dziedzic, Optical trapping and manipulation of viruses and bacteria. Science, 1987. 235(4795): p. 1517–1520.

34. Ashkin, A., J.M. Dziedzic, and T. Yamane, Optical trapping and manipulation of single cells using infrared laser beams. Nature, 1987. 330(6150): p. 769–771.

35. Kong, L., et al., Characterization of bacterial spore germination using phase-contrast and fluorescence microscopy, Raman spectroscopy and optical tweezers. Nature Protocols, 2011. 6(5): p. 625.

36. MacDonald, M.P., G.C. Spalding, and K. Dholakia, Microfluidic sorting in an optical lattice. Nature, 2003. 426(6965): p. 421–424.

37. Paterson, L., et al., Controlled rotation of optically trapped microscopic particles. Science, 2001. 292(5518): p. 912–914.

38. Thalhammer, G., et al., Optical macro-tweezers: trapping of highly motile micro-organisms. Journal of Optics, 2011. 13(4): p. 044024.

39. Liphardt, J., et al., Equilibrium information from nonequilibrium measurements in an experimental test of Jarzynski’s equality. Science, 2002. 296(5574): p. 1832–1835.

40. delToro, D. and D.E. Smith, Accurate measurement of force and displacement with optical tweezers using DNA molecules as metrology standards. Applied Physics Letters, 2014. 104(14): p. 143701.

41. Keyser, U., et al., Optical tweezers for force measurements on DNA in nanopores. Review of Scientific Instruments, 2006. 77(10): p. 105105.

42. Wang, C., et al., Photophoretic trapping-Raman spectroscopy for single pollens and fungal spores trapped in air. Journal of Quantitative Spectroscopy and Radiative Transfer, 2015. 153: p. 4–12.

43. Rouger, V., et al., Independent synchronized control and visualization of interactions between living cells and organisms. Biophysical Journal, 2014. 106(10): p. 2096–2104.

44. Pham, V.H. and J. Kim, Cultivation of unculturable soil bacteria. Trends in Biotechnology, 2012. 30(9): p. 475–484.

45. Hawes, C., et al., Optical tweezers for the micromanipulation of plant cytoplasm and organelles. Current Opinion in Plant Biology, 2010. 13(6): p. 731–735.

46. Greulich, K., et al., Micromanipulation by laser microbeam and optical tweezers: from plant cells to single molecules. Journal of Microscopy, 2000. 198(3): p. 182–187.

47. Gao, H., et al., In vivo quantification of peroxisome tethering to chloroplasts in tobacco epidermal cells using optical tweezers. Plant Physiology, 2016. 170(1): p. 263–272.

48. Abe, Y., et al., Micromanipulation of amyloplasts with optical tweezers in Arabidopsis stems. Plant Biotechnology, 2020: p. 20.1201 a.

49. Christensen, B., Physical fractionation of soil and structural and functional complexity in organic matter turnover. European Journal of Soil Science, 2001. 52(3): p. 345–353.

50. Sessitsch, A., et al., Microbial population structures in soil particle size fractions of a long-term fertilizer field experiment. Applied and Environmental Microbiology, 2001. 67(9): p. 4215–4224.

51. Ciani, A., K.U. Goss, and R.P. Schwarzenbach, Light penetration in soil and particulate minerals. European Journal of Soil Science, 2005. 56(5): p. 561–574.

52. Or, D., et al., Physical constraints affecting bacterial habitats and activity in unsaturated porous media–a review. Advances in Water Resources, 2007. 30(6-7): p. 1505–1527.

53. Roichman, Y., et al., Optical traps with geometric aberrations. Applied Optics, 2006. 45(15): p. 3425–3429.

54. Tam, J.M., I. Biran, and D.R. Walt, Parallel microparticle manipulation using an imaging fiber-bundle-based optical tweezer array and a digital micromirror device. Applied Physics Letters, 2006. 89(19): p. 194101.

55. Čižmár, T., M. Mazilu, and K. Dholakia, In situ wavefront correction and its application to micromanipulation. Nature Photonics, 2010. 4(6): p. 388–394.

56. McGloin, D. and K. Dholakia, Bessel beams: diffraction in a new light. Contemporary Physics, 2005. 46(1): p. 15–28.

57. Ng, J., Z. Lin, and C. Chan, Theory of optical trapping by an optical vortex beam. Physical Review Letters, 2010. 104(10): p. 103601.

58. Zhao, Z., W. Zang, and J. Tian, Optical trapping and manipulation of Mie particles with Airy beam. Journal of Optics, 2016. 18(2): p. 025607.

59. Gong, L., et al., Generation of cylindrically polarized vector vortex beams with digital micromirror device. Journal of Applied Physics, 2014. 116(18): p. 183105.

60. Suarez, R.A., et al., Experimental optical trapping with frozen waves. Optics Letters, 2020. 45(9): p. 2514–2517.

61. Thalhammer, G., et al., Acoustic force mapping in a hybrid acoustic-optical micromanipulation device supporting high resolution optical imaging. Lab on a Chip, 2016. 16(8): p. 1523–1532.

62. De Vlaminck, I. and C. Dekker, Recent advances in magnetic tweezers. Annual Review of Biophysics, 2012. 41: p. 453–472.

63. Dholakia, K., B.W. Drinkwater, and M. Ritsch-Marte, Comparing acoustic and optical forces for biomedical research. Nature Reviews Physics, 2020. 2(9): p. 480–491.

64. Neuman, K.C. and A. Nagy, Single-molecule force spectroscopy: optical tweezers, magnetic tweezers and atomic force microscopy. Nature Methods, 2008. 5(6): p. 491–505.

65. Hall-Stoodley, L., J.W. Costerton, and P. Stoodley, Bacterial biofilms: from the natural environment to infectious diseases. Nature Reviews Microbiology, 2004. 2(2): p. 95–108.

66. Bialek, W. and S. Setayeshgar, Physical limits to biochemical signaling. Proceedings of the National Academy of Sciences, 2005. 102(29): p. 10040–10045.

67. Li, G., L.-K. Tam, and J.X. Tang, Amplified effect of Brownian motion in bacterial near-surface swimming. Proceedings of the National Academy of Sciences, 2008. 105(47): p. 18355–18359.

68. Chenu, C., Interactions between microorganisms and soil particles: an overview. Interactions between soil particles and microorganisms: Impact on the Terrestrial Ecosystem, 2002. 1: p. 1–40.

69. Chia, T.W.R., et al., Stochasticity of bacterial attachment and its predictability by the extended Derjaguin-Landau-Verwey-Overbeek theory. Applied and Environmental Microbiology, 2011. 77(11): p. 3757–3764.

70. Aung, K., Y. Jiang, and S.Y. He, The role of water in plant–microbe interactions. The Plant Journal, 2018. 93(4): p. 771–780.

71. Bogino, P.C., et al., The role of bacterial biofilms and surface components in plant-bacterial associations. International Journal of Molecular Sciences, 2013. 14(8): p. 15838–15859.

72. López, H.M., et al., Turning bacteria suspensions into superfluids. Physical Review Letters, 2015. 115(2): p. 028301.

73. Guber, A., D. Shelton, and Y.A. Pachepsky, Effect of manure on Escherichia coli attachment to soil. Journal of Environmental Quality, 2005. 34(6): p. 2086–2090.

74. Nadarasah, G. and J. Stavrinides, Insects as alternative hosts for phytopathogenic bacteria. FEMS Microbiology Reviews, 2011. 35(3): p. 555–575.

75. Aira, M., M. Pérez-Losada, and J. Domínguez, Diversity, structure and sources of bacterial communities in earthworm cocoons. Scientific Reports, 2018. 8(1): p. 1–9.

